# Context-Aware Synthetic Promoter Design Using Neural Networks Enables Rewiring of Eukaryotic Transcriptional Networks

**DOI:** 10.1101/2025.08.27.672570

**Authors:** L Kuhajda, T Honzík, J Švec, D Georgiev

## Abstract

Gene regulation through promoter engineering is a cornerstone of synthetic biology, enabling precise control over transcriptional networks. However, experimental approaches remain labor-intensive. While artificial neural networks (ANNs) have improved regulatory element prediction, tools for promoter–transcription factor binding site (TFBS) recombination are still lacking. We present an ANN framework for context-aware design of synthetic promoters in *Saccharomyces cerevisiae*. The model predicts optimal TFBS insertion sites and the extent of promoter rewriting needed for successful integration. Applying this, we screened 6,011 native yeast promoters for compatibility with the TetR TFBS, generating a ranked list of high-confidence promoter–TFBS pairs. Experimental validation showed that model-designed promoters achieved repression rates up to 98.4%, without prior experimental characterization or tuning. We further rewired the yeast transcriptional network by introducing glucose-dependent regulation of an essential gene via Mig1 TFBS insertion. These results establish a scalable, predictive method for engineering regulatory sequences and reprogramming transcriptional logic.

## Introduction

Cells regulate gene expression through complex networks of proteins, RNA molecules, and DNA sequences [1]. These regulatory systems allow organisms to respond to environmental changes, coordinate functions, and develop adaptive or novel traits. Rewiring these transcriptional networks by modifying regulatory elements can enable entirely new cellular behaviors and is a key goal in synthetic biology [2, 3, 4].

Rewiring of transcriptional networks largely relies on the use of synthetic promoter libraries that lack the context-dependence of native promoters. Over the years, various synthetic promoter libraries have been developed to control expression strength or make promoters responsive to specific inputs by inserting transcription factor binding sites (TFBSs) [5-11]. These engineered promoters have been applied in metabolic engineering [5, 12-14], biosensors, and regulatory devices like toggle switches, logic gates, and oscillators [14-18]. Native promoters exhibit context-dependent behavior influenced by environmental conditions and genetic background. This sensitivity is retained when promoters serve as scaffolds for TFBS insertions, allowing for the construction of dynamically regulated synthetic promoters [19-23].

Despite advances in DNA editing and assembly [24-26], promoter engineering remains difficult due to the lack of conserved structures to guide insertion and deletion of regulatory sequences. Artificial neural networks (ANNs) have shown strong potential for resolving regulatory sequences and guiding design. Several models have been used to predict promoter strength, enhancer activity, or chromatin accessibility (Basset [27], DanQ [28], Basenji [29]). Other models focus on enhancer-promoter interactions (SPEID [30], DeepSEM [31], Enformer [32]) or a promoter strength prediction from experimental data with subsequent generative models to design promoters with specific strength [33-39]. However, current tools do not predict potential recombinations of promoters and TFBSs, a critical step for building context-aware, responsive regulatory elements.

We developed a two-stage ANN system that uses context-aware sequence recombination to propose general TFBS insertion. The first model (*Place-Back*) identifies the most suitable insertion region, and the second model (*Determiner*) refines the predicted rewrite length to ensure a good fit. The models are trained using a self-supervised approach, allowing them to learn the context of TFBS insertions directly from genomic data without any labels.

We applied the system to screen 6,011 native yeast promoters for compatibility with the TetR TFBS (tetO), generating a ranked list of high-confidence promoter-TFBS pairs.

In addition, we validated two unique promoter systems *in vivo*: (1) untested wildtype *S*.*cerevisiae* promoters were regulated by the TetR repressing system, achieving strong conditional repression up to 98.4% with no prior experimental characterization and tuning, and (2) the *S*.*cerevisiae* wildtype transcriptional regulatory network was rewired to create a completely new glucose sensitive response achieved through glucose-responsive regulation of an essential gene. Together, these results show that context-aware promoter design using neural networks can streamline synthetic promoter construction and enable more complex regulatory systems.

The developed tool is available as an online browser-based application and as a locally executable version via the associated GitHub repository. Access details are provided in the “Code Availability” section.

## Results

### Dataset

The target organism for this study is *Saccharomyces cerevisiae*. To train an artificial neural network with evolutionarily relevant sequences, we extracted promoter sequences for all annotated genes from 25 *Saccharomycotina* species. These sequences were split into training and validation datasets, while *S*.*cerevisiae* promoters were excluded and reserved solely for testing. Promoters were defined as 400 bp upstream of the respective start codon, ensuring at least 100bp separation from the nearest neighboring gene in the promoter direction. This resulted in a dataset of 6,011 *S*.*cerevisiae* promoters for testing, and 133,889 promoters from *Saccharomycotina* further divided into training dataset with 123,889 promoters, and validation dataset with 10,000 promoters. All sequences were stored in nucleotide form for model training and evaluation.

### Two-Stage Model Architecture for TFBS Insertion

To support context-aware promoter engineering, we built a two-stage artificial neural network (ANN) system made up of two models. The first model, called *Place-Back*, predicts the most suitable site for inserting a transcription factor binding site (TFBS) into a given promoter sequence with two variants trained, differing in the *Model-Specific Rewriting Length* (MSRL): one replaces 5 bp (closely modeling actual TFBS insertion) and the other 40 bp (modeling larger-scale sequence replacement). The second model, called *Determiner*, then decides how much of the promoter should be overwritten, either a short (5 bp) or long (40 bp) region, and identifies the most likely region of insertion. The two models are described in detail below.

The *Place-Back* model uses a self-supervised training approach. To prepare a training sample (Fig. 1A.1), two promoters (400bp each) from the same yeast species are randomly selected: *base-promoter* and *auxiliary-promoter*. A segment of 8-64 bp is extracted from the *base-promoter* to serve as the query sequence, and this query is replaced in its original position by a MSRL-length segment from the *auxiliary-promoter*. To prevent the model from simply detecting the query region by alignment, we also introduce at random positions 2-3 additional disruptions (segments of MSRL length) to the *base-promoter* from the *auxiliary-promoter*. This forces the model to learn contextual features rather than relying on exact sequence matching. *Place-Back* model takes the disrupted *base-promoter* and query as input and outputs a 400-length prediction vector indicating the probability that each base is part of the original query region. The model consists of embedding layer with positional encoding for nucleotide representation, convolutional neural networks (CNNs) to reduce promoter length and capture local motifs, a transformer encoder to learn long-range dependencies, a bi-directional LSTM to process query sequence, dot-product based attention mechanism to combine both inputs, and final sigmoid activation to generate base-pair level probabilities (Fig. 1A.2,3). Two *Place-Back* models with MSRL of 5bp and two models with MSRL of 40bp were trained. Each model contains ∼6 million parameters and was trained for 168 hours on a GeForce GTX 1080 Ti GPU, processing roughly 300 million training examples.

**Figure 1:**
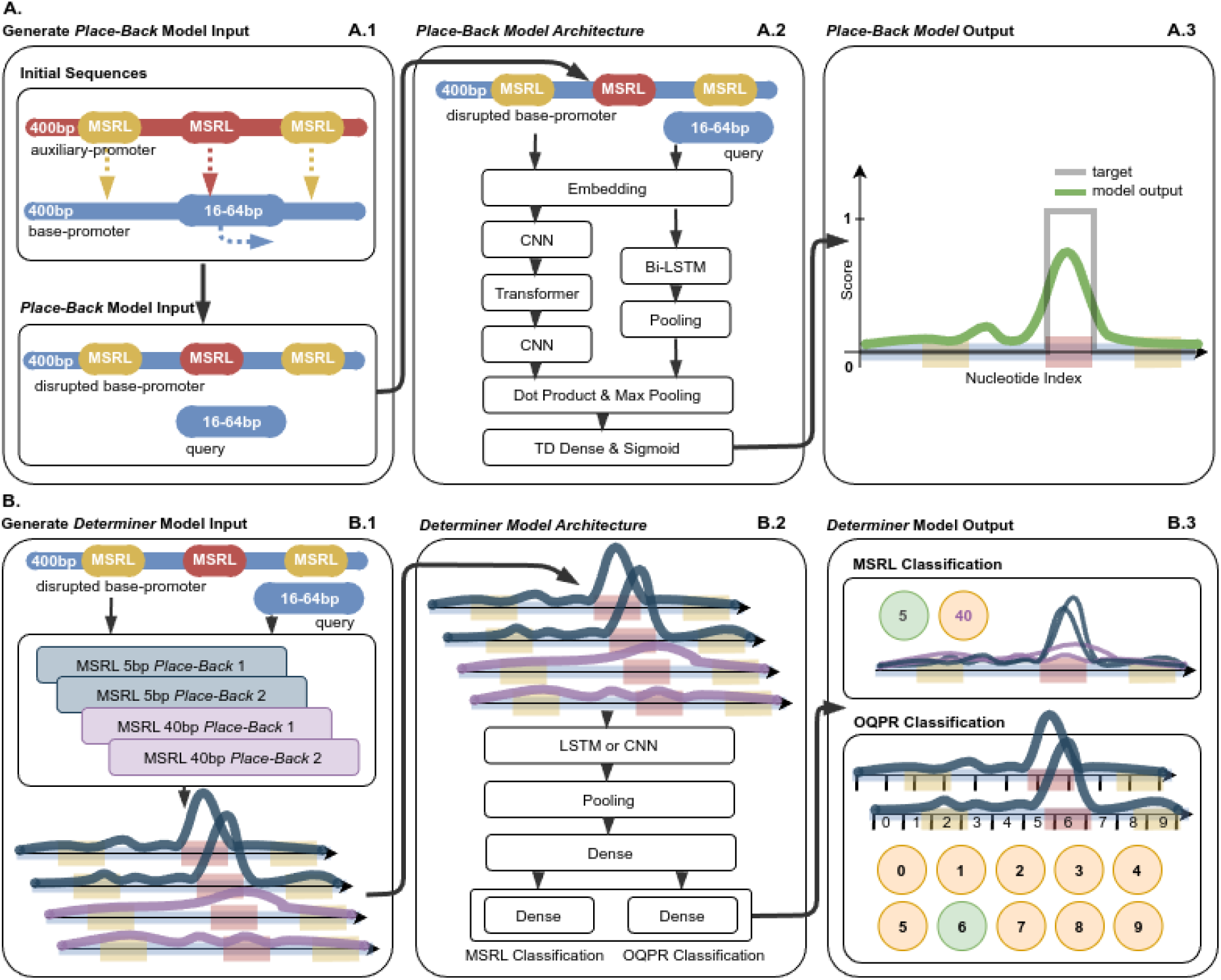
Architecture and training of the two-stage ANN system for TFBS insertion. (A.1) **Training data generation for Place-Back models:** Two 400 bp promoter sequences are randomly selected, *base-promoter* and *auxiliary-promoter*. A 16–64 bp segment is extracted from the *base-promoter* (query) and replaced with a 5 bp or 40 bp (MSRL) segment from the other (*auxiliary-promoter*). To prevent trivial learning, 2–3 additional disruptions of fixed length (MSRL) are introduced at random positions. (A.2) **Place-Back model architecture:** The disrupted *base-promoter* is processed through embedding, CNNs for compression, a transformer layer, and CNNs for decompression. The query is processed via embedding and Bi-LSTM. The two branches are combined with a dot-product operation, followed by max pooling and a sigmoid output over all 400 promoter positions. (A.3) **Model output:** The resulting prediction is a vector of probabilities (0–1) across the promoter, with peaks indicating the predicted probability of the original query location within the *base-promoter*. The model is trained to assign values approaching 1 at the query’s original location and 0 elsewhere. (B.1) **Input to Determiner models:** A single disrupted *base-promoter* and query sequence pair is passed through four *Place-Back* models (two for MSRL=5bp, two for MSRL=40bp), generating four prediction curves. (B.2) **Determiner model architecture:** The stacked *Place-Back* outputs are processed through a CNN or LSTM-based model, followed by dense layers. (B.3) **Determiner output:** The model predicts (1) the correct rewrite length (5bp or 40bp) and (2) the most likely insertion region (divided into 10 bins of 40 bp each).

Outputs from the *Place-Back* models are passed to the *Determiner* model (Fig. 1B.1), which resolves inconsistencies between predictions and classifies the most likely insertion region and required rewrite length. For training, the *Determiner* receives the output curves from two 5bp and two 40 bp *Place-Back* models for a single input sample generated with either MSRL of 5 bp or 40 bp. The *Determiner* predicts the rewrite length the sample was generated with, and the *Original Query Position Region* (OQPR) - one of ten 40 bp regions spanning the promoter (Fig. 1B.3). Ten *Determiner* models with diverse CNN or LSTM architectures (1.8-13.6 million parameters) were trained. Consensus across models was used to increase robustness: only predictions where at least 7/10 *Determiner* models agreed were considered high-confidence.

### Model Performance and Prediction Filtering

Model performance was evaluated on held-out validation data. The *Place-Back* models reached an average precision (accuracy of positive predictions) of 48%, meaning that nearly half of the positive predictions (top-scoring insertion locations) matched the original query placement. Recall (coverage of actual positives) values depended on the MSRL: 5 bp models had lower recall (∼4%) due to narrower and lower prediction peak values, while 40 bp models had higher recall (∼32%).

While these precision and recall values may appear modest by conventional machine learning standards, they are both statistically meaningful and biologically actionable in this context. The task involves locating a functional insertion site within a 400 bp promoter sequence, an inherently difficult and imbalanced classification problem. Even moderate precision therefore represents a significant enrichment over random chance. Importantly, promoter functionality in vivo is often tolerant to slight positional variation, meaning that predictions considered incorrect by strict matching criteria may still result in successful TFBS integration. This is supported by the strong experimental performance of selected model predictions, described in the sections below.

To further increase the reliability of predictions, we introduced a filtering step based on multi-model consensus using the *Determiner*. Only predictions with strong internal consistency across multiple models were retained. This strategy provides a practical interface for experimental selection.

### In Silico Recombination of Yeast Promoters with tetO

In an inference mode the TFBS with length 8-64bp enters the system as the query sequence, the promoter enters in wild-type form. An example of the overall output of the model for recombination of a wild-type promoter with a TFBS is shown in Fig. 2.

**Figure 2:**
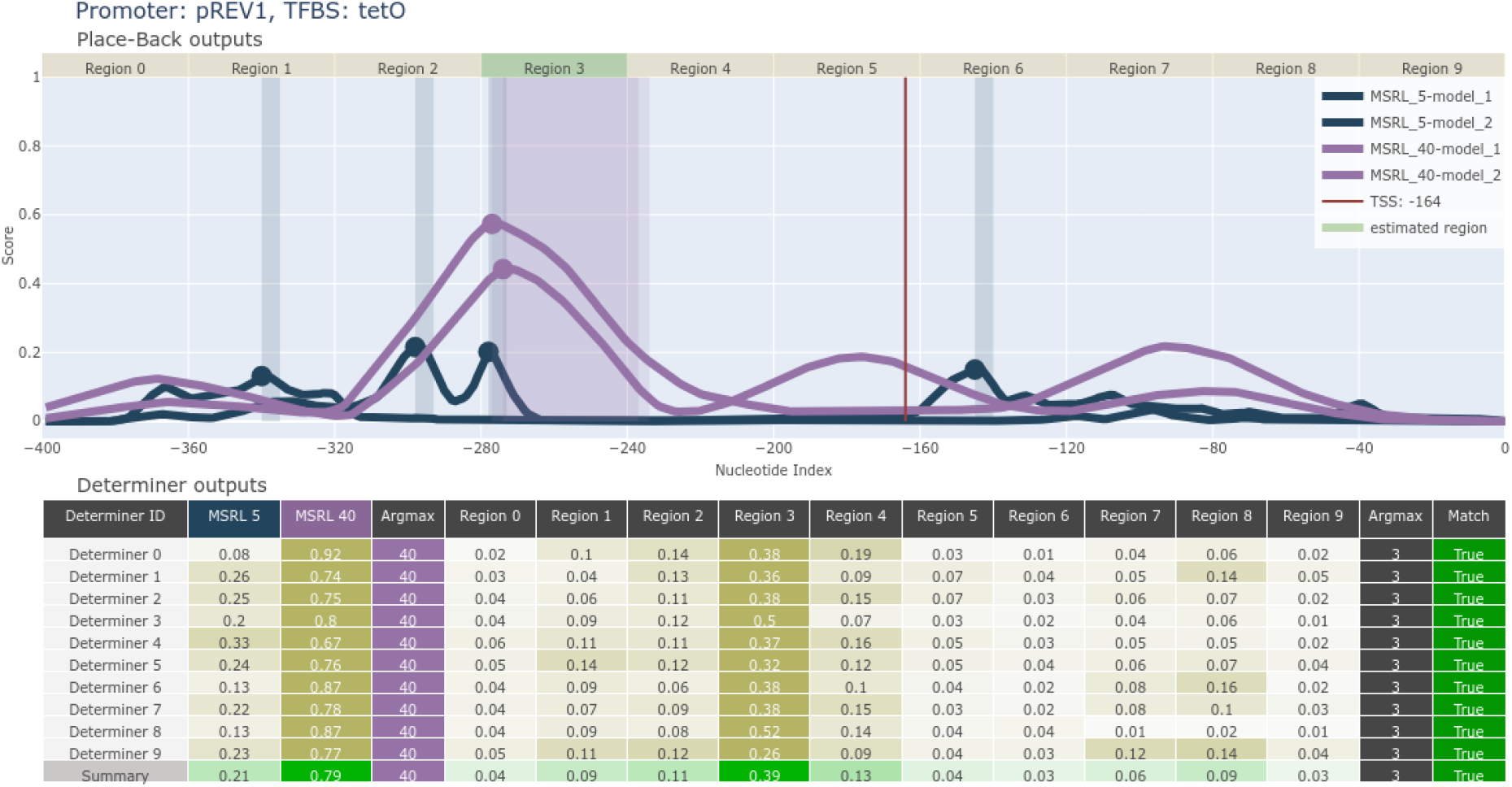
Example output from the two-stage ANN system for tetO recombination into the pREV1 promoter. **Top panel**: Outputs from four *Place-Back* models. Two 5bp models (blue) and two 40bp models (purple) show insertion probability across the promoter sequence. The red line marks the transcription start site (TSS). **Bottom panel**: Predictions from ten *Determiner* models. All *Determiner* models select MSRL=40bp and region 3 as the optimal insertion window. The highlighted region in the top panel corresponds to this classification, confirming a high-confidence recombination proposal. Based on the outputs, tetO would be rewritten to the promoter region from −277 to −237.

To evaluate the potential of the model in a real use case, we applied it to all 6,011 *S*.*cerevisiae* promoters (defined as 400 bp upstream of the start codon). Each promoter was virtually recombined with the TetR TFBS (tetO: TCCCTATCAGTGATAGAGATCTCCCTATCAGTGATAGAGA [40]). The model evaluated whether tetO could be inserted into each promoter without disrupting essential elements like the TATA-box or transcription start site (TSS). Of the 6,011 promoters, 3,712 were recommended for recombination, while 2,299 were filtered out due to a lack of model consensus or predicted disruption of minimal regulatory elements.

Among the recommended promoters, 2,149 were assigned to use a 5bp rewrite and 1,560 were assigned to use a 40bp rewrite. Analysis of the predicted insertion sites showed that, for 5 bp modifications, 85% occurred in the distal half of the promoter, and 60% were within the last 100 bp upstream of the start codon (Supplementary Fig. 1). For 40 bp rewrites, about 50% of insertions also occurred in this downstream region (Supplementary Fig. 4). When TATA-box or TSS positions were known, a notable fraction of insertions occurred within 30 bp of the TATA-box (25%) or within 33 bp of the TSS (20%) (Supplementary Fig. 5,6). These patterns suggest the model prefers insertions near core regulatory regions but avoids direct disruption. A full list of promoter-TFBS pairs and insertion details is available in the Supplementary Materials.

### Experimental Validation of Synthetic Promoters with TetR

To experimentally validate the ANN predictions, we selected four *S*.*cerevisiae* constitutive promoters of different strengths (pHHF2, pPAB1, pPOP6, pREV1) predicted to be compatible with tetO insertion, for experimental testing. Each promoter was recombined with tetO at the position suggested by the model, either 5bp or 40bp rewrite (system output with position selection shown in Supplementary Fig. 7-10), and fused to a NanoLuc reporter.

This system was orthogonal: the TetR repressor was introduced on a separate expression cassette (pCCW12-TetR), and TetR is not natively present in yeast. This means the repression system could be tested without interference from endogenous transcription factors, allowing clean validation of the ANN-designed promoter-TFBS combinations.

The synthetic promoters were tested in strains with and without TetR expression, with and without doxycycline induction to set deactivation of repression (Fig. 3.A). All four synthetic promoters showed strong repression in the OFF state (TetR expressed, no doxycycline), confirming the function of the model-designed recombination. Repression rates were: pPOP6_tetO - 98.4% (64.1-fold induction), pPAB1_tetO - 90.1% (10.2-fold), pREV1_tetO - 62.7% (2.7-fold), and pHHF2_tetO - 35.8% (1.6-fold) (Fig. 3.B,C). These results demonstrate that the model can identify insertion sites that enable high dynamic range regulation without manual tuning.

**Figure 3:**
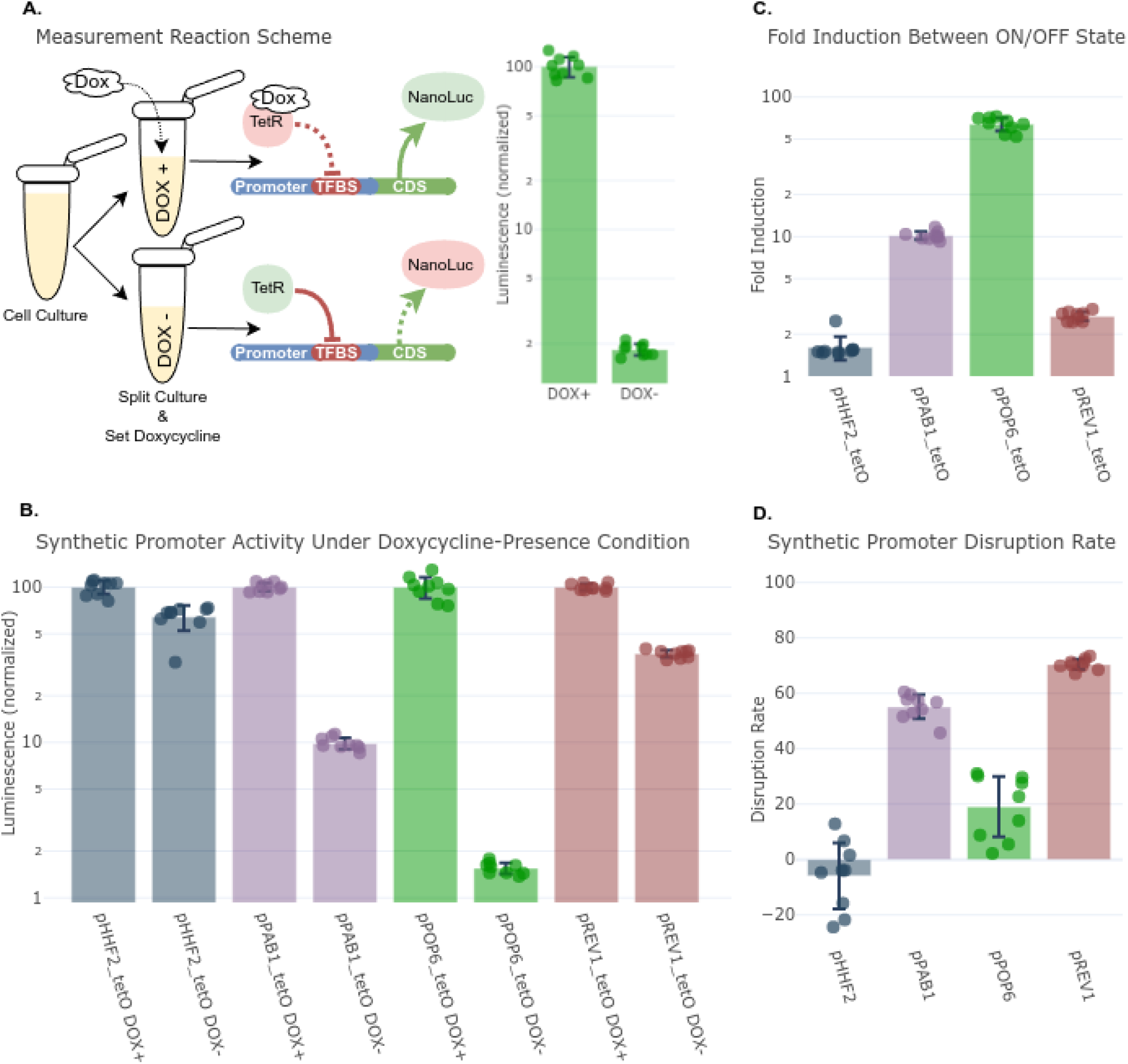
Experimental validation of ANN-designed synthetic promoters recombined with tetO and tested using an orthogonal TetR repression system. **Experimental setup**: Synthetic promoters were fused to NanoLuc and integrated into yeast strains with or without TetR expression. Promoter activity was measured in ON (doxycycline present) and OFF (TetR active) states. **Repression levels** of each synthetic promoter normalized to ON-state activity (100%). pPOP6_tetO showed the strongest repression (98.4%), followed by pPAB1_tetO (90.1%), pREV1_tetO (62.7%), and pHHF2_tetO (35.8%). **Fold induction**: pPOP6_tetO exhibited a 64-fold ON/OFF induction, followed by pPAB1_tetO (10.2-fold), pREV1_tetO (2.7-fold), and pHHF2_tetO (1.6-fold). **Impact on native activity**: Comparison of wild-type vs. synthetic promoter activity in the absence of TetR. pHHF2 showed no disruption, pPOP6 activity dropped by 19%, while pPAB1 and pREV1 exhibited 55% and 70% activity loss, respectively.

We also evaluated how tetO insertion affected native promoter strength by comparing wild-type vs. synthetic versions in strains lacking TetR (Fig. 3.D). Some promoters retained most of their activity (pHHF2 was unaffected, pPOP6 dropped 19%), while others showed moderate to stronger reduction (pPAB1 dropped 55%, pREV1 dropped 70%).

These results confirm that ANN-based design can yield strong, tunable repression, and that the models can correctly identify sites with minimal disruption to native promoter activity. The orthogonal design enabled clean evaluation of promoter behavior in isolation.

### Rewiring of the Yeast Transcriptional Network

To demonstrate that the system can be used not only for orthogonal control but also for rewiring native transcriptional regulation, we engineered the promoter of PCF11, an essential gene [41], by inserting a Mig1 repressor binding site (mig1O: GTATTAAACCCGGGGTA [42]).

Mig1 is a glucose-sensitive transcriptional repressor active in high-glucose conditions (Fig. 4.A) [43-45]. Using the model, we inserted mig1O in the PCF11 promoter (Supplementary Fig. 11). Unlike the TetR system, this construct relied entirely on endogenous regulation, so no extra TFs were introduced.

**Figure 4:**
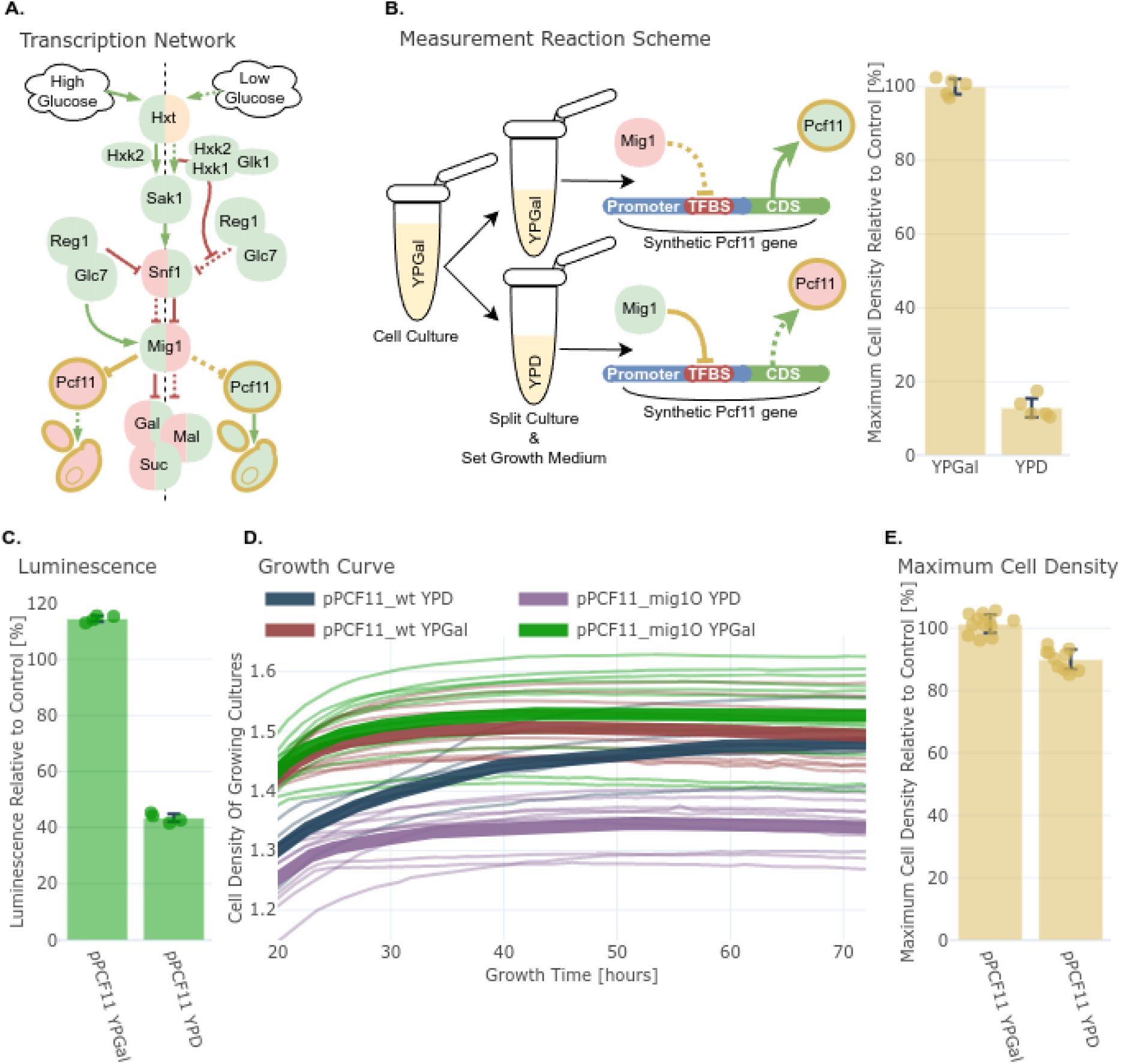
ANN-guided rewiring, insertion of Mig1 TFBS into the essential PCF11 promoter enables glucose-dependent transcriptional repression and growth control. **(A) Schematic**: The PCF11 promoter was modified with a Mig1 binding site (mig1O), creating a new regulatory edge in the glucose repression network. **(B) Experimental workflow**: Strains with synthetic and wild-type PCF11 promoters were tested in glucose (Mig1 ON) and galactose (Mig1 OFF). **(C) Luciferase assay**: In glucose, pPCF11_mig1O showed 57% repression compared to wild-type; in galactose, expression was slightly higher than wild-type. **(D) Growth curves**: Strains with the synthetic promoter showed 10% reduced maximum cell density in glucose and 1.4% increased in galactose, consistent with conditional repression of an essential gene. **(E) Growth summary**: Bar plot showing the relative change in maximal cell density. The results confirm successful network rewiring with regulatory effects that are strong but not lethal.

First, we validated glucose-dependent regulation using NanoLuc assays. In galactose (Mig1 OFF, Fig. 4.B), the synthetic promoter showed 14% stronger activity than wild-type (Fig. 4.C). In glucose (Mig1 ON, Fig. 4.B), it was repressed by 57%.

Next, we replaced the endogenous PCF11 promoter in the yeast genome with the synthetic pPCF11_mig1O promoter using CRISPR-Cas9. Because PCF11 is essential, full repression would be expected to be lethal. Growth assays showed that in galactose, the strain with pPCF11_mig1O had slightly higher growth than wild-type (+1.4%) and in glucose, where pPCF11_mig1O is repressed by active Mig1, the strain had a 10% lower maximum cell density than wild-type, confirming repression but not full loss of function (Fig. 4.D,E).

This result is a key proof-of-concept: a single ANN-designed TFBS insertion created a new regulatory edge to rewire the yeast transcription network, controlling expression of an essential gene based on nutrient conditions, without additional synthetic TFs.

## Discussion

We developed a two-stage artificial neural network system capable of guiding transcription factor binding site (TFBS) insertion into native promoter sequences in a context-aware manner. By combining self-supervised learning with promoter architecture analysis, our framework identifies viable insertion sites and proposes minimal, functional sequence modifications. The model was validated both in silico and in vivo, successfully generating synthetic promoters with strong repression in an orthogonal TetR system and enabling conditional control of an essential gene through rewiring of the native transcriptional network.

In the first phase, the implementation of an orthogonal repressor system using TetR-responsive promoters successfully introduced tunable transcriptional regulation into yeast. The designed synthetic promoters showed varying degrees of repression, with pPOP6_tetO achieving the highest repression rate of 98.4% and pPAB1_tetO reaching 90.1%. The analysis of promoter activity in wild-type versus engineered constructs indicated that while some promoters retained their original strength, others exhibited disruptions due to the introduction of transcription factor binding sites (TFBS). Despite these variations, the system effectively enabled external regulation of synthetic gene expression, demonstrating the power of ANN-driven promoter engineering.

The second phase of the study expanded upon these results by introducing a new regulatory edge within the native transcription network of *S*.*cerevisiae*. The pPCF11 promoter corresponding to the essential gene was engineered with a Mig1 repressor binding site, allowing for conditional repression based on glucose availability. Luciferase assays confirmed that the synthetic promoter responded to glucose-mediated repression, with a 57% repression rate in YPD medium. When the rewired regulatory system was implemented in the yeast genome, growth curve analysis revealed a 10% reduction in maximal cell density under high-glucose conditions, confirming that the introduced repression was functional but not lethal.

Overall, this study highlights the potential of ANN design in synthetic biology, enabling precise genetic modifications while minimizing unintended cellular disruptions. The success of both the orthogonal gene control and transcription network rewiring demonstrates that data-driven approaches can be leveraged for predictive and efficient genetic engineering.

Future work will focus on enhancing the ANN models for their applicability beyond *S*.*cerevisiae* to include other eukaryotic organisms such as plants, invertebrates, mammals and other vertebrates.

## Methods

### Strains and Plasmids Preparation

All experiments were performed in *Saccharomyces cerevisiae*. The yeast strain BY4741 was used as the parental strain (see Table 1 for full genotypes). To create a strain expressing the TetR repressor, a cassette containing pCCW12-TetR and a LEU2 selection marker was integrated into BY4741, resulting in strain S100. A control strain (S101) was constructed by integrating an empty LEU2 cassette.

**TABLE 1:** strains.

| strain | genotype | reference |
| --- | --- | --- |
| BY4741 | MATa his3Δ1 leu2Δ0 met15Δ0 ura3Δ0 | [49] |
| S100 | BY4741 leu2::pCCW12-TetR |  |
| S101 | BY4741 leu2::empty cassette |  |
| S102 | BY4741 leu2::pCCW12-TetR his3::pHHF2_wt-NanoLuc |  |
| S103 | BY4741 leu2::empty cassette his3::pHHF2_wt-NanoLuc |  |
| S104 | BY4741 leu2::pCCW12-TetR his3::pHHF2_tetO-NanoLuc |  |
| S105 | BY4741 leu2::empty cassette his3::pHHF2_tetO-NanoLuc |  |
| S106 | BY4741 leu2::pCCW12-TetR his3::pPAB1_wt-NanoLuc |  |
| S107 | BY4741 leu2::empty cassette his3::pPAB1_wt-NanoLuc |  |
| S108 | BY4741 leu2::pCCW12-TetR his3::pPAB1_tetO-NanoLuc |  |
| S109 | BY4741 leu2::empty cassette his3::pPAB1_tetO-NanoLuc |  |
| S110 | BY4741 leu2::pCCW12-TetR his3::pPOP6_wt-NanoLuc |  |
| S111 | BY4741 leu2::empty cassette his3::pPOP6_wt-NanoLuc |  |
| S112 | BY4741 leu2::pCCW12-TetR his3::pPOP6_tetO-NanoLuc |  |
| S113 | BY4741 leu2::empty cassette his3::pPOP6_tetO-NanoLuc |  |
| S114 | BY4741 leu2::pCCW12-TetR his3::pREV1_wt-NanoLuc |  |
| S115 | BY4741 leu2::empty cassette his3::pREV1_wt-NanoLuc |  |
| S117 | BY4741 leu2::pCCW12-TetR his3::pREV1_tetO-NanoLuc |  |
| S118 | BY4741 leu2::empty cassette his3::pREV1_tetO-NanoLuc |  |
| S200 | BY4741 his3::pPCF11_wt-NanoLuc |  |
| S201 | BY4741 his3::pPCF11_mig1O-NanoLuc |  |
| S300 | BY4741 pPCF11::pPCF11_mig1O |  |

Promoters selected for testing (pHHF2, pPAB1, pPOP6, pREV1) were obtained as domesticated part plasmids from the MoClo Yeast Toolkit [46] and represented a range of constitutive expression strengths. These promoters are referred to as pX_wt, where “X” is the gene source, and “wt” indicates wild-type sequence.

### Construction of Synthetic Promoters with TetR Binding Sites

To insert TetR operator sequences (tetO) into native promoters, we used a scarless split-PCR method guided by the ANN model predictions. Each promoter was split at a model-predicted insertion point with a gap based on selected MSRL of 5bp or 40 bp into two fragments: pX_frag1 (upstream of insertion) and pX_frag2 (downstream of insertion). Restriction sites were strategically placed to allow seamless assembly of the fragments flanking the inserted tetO sequence, avoiding any extraneous nucleotides at junctions.

- pX_frag1 included a 5′ overhang with the upstream portion of tetO and a BsmBI site for joining with pX_frag2, and a 3′ BsaI/BsmBI site for cloning.
- pX_frag2 contained the complementary BsmBI site and the downstream portion of tetO in its 3′ overhang, and a 5′ BsaI/BsmBI site

The two promoter fragments with tetO insert were assembled via BsmBI Golden Gate assembly into a CamR-marked bacterial entry vector (PEV), producing synthetic promoter constructs referred to as pX_tetO, where “tetO” refers to promoter version recombined with TFBS tetO. Assembly products were transformed into *E*.*coli* NEB Turbo cells and screened via colony PCR and sequencing.

### Assembly of Expression Constructs

Final expression cassettes were assembled using BsaI Golden Gate reactions. Each construct contained:

- A synthetic or wild-type promoter (pX_tetO or pX_wt),
- His3 selection marker (with AmpR for E. coli propagation),
- MFW PrePro signal peptide,
- NanoLuc reporter [47],
- tENO2 terminator.

Following transformation into NEB Turbo *E*.*coli*, plasmid miniprep was performed, and constructs were validated by BsmBI restriction digestion and gel electrophoresis.

### Genomic Integration into Saccharomyces cerevisiae

Final gene constructs were integrated into *S*.*cerevisiae* strains S100 and S101 at designated orthogonal loci via yeast transformation and plated on His3 selection media, resulting in strains S102-S118. Successful genomic integration was confirmed through NotI digestion analysis and colony PCR.

### Luciferase Assays for Promoter Rewiring Experiments

To assess transcriptional rewiring, the native pPCF11 promoter (750 bp upstream of the ORF) was amplified and modified by insertion of a Mig1 operator sequence (mig1O), using the same scarless split-PCR strategy described above. The resulting synthetic promoter (pPCF11_mig1O) and wild-type version (pPCF11_wt) were assembled with NanoLuc, HIS3, and tENO2, then integrated into BY4741, producing strains S200 and S201 for luciferase-based expression analysis.

### CRISPR-Cas9 Mediated Genomic Rewiring

Precise genomic integration of pPCF11_mig1O was performed using CRISPR-Cas9. A guide RNA targeting a PAM-adjacent site in the native pPCF11 promoter was cloned into a Cas9 expression plasmid with a *URA3* marker. The synthetic pPCF11_mig1O construct was supplied as a donor repair template. Co-transformation into BY4741 yielded strain S300. Transformants were selected on *URA3*-deficient media, and the Cas9 plasmid was subsequently removed via 5-FOA counter-selection. Correct integration was confirmed by PCR and sequencing.

### Gene Expression Analysis Using Luciferase Assays

Promoter activity was measured using NanoLuc luciferase assays under different environmental conditions.

TetO promoters: Cultures were grown in YPD media [48], diluted to OD600 = 0.05, and treated with 2 μg/mL doxycycline (DOX) (ON state) or left untreated (OFF state). Relative Light Unit (RLU) was measured after 2 hours in a white 96-well plate. Promoter activity and regulatory metrics were calculated as follows:

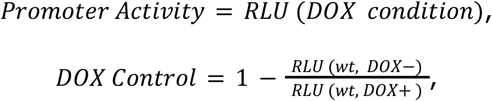

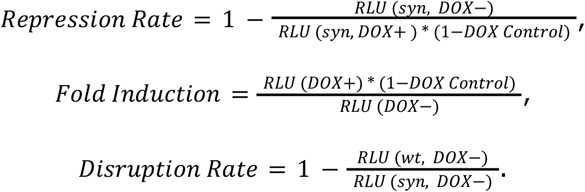

Mig1O promoters: Strains were cultured in YPGal media (YPD [48] with glucose replaced by the same concentration of galactose), diluted to OD600 = 0.05, and split into YPD (high glucose, repression ON) and YPGal (low glucose, repression OFF). Relative promoter activity was determined as:

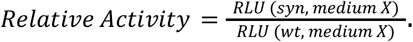

### Maximum Cell Density Measurements for Rewired Network

To evaluate the effect of pPCF11_mig1O on growth, strains BY4741 (wild-type pPCF11_wt in the genome) and S300 (synthetic promoter pPCF11_mig1O in the genome) were cultured in YPGal, and washed to YPD and YPGal media. Cultures were diluted to OD600 = 0.05 and monitored every hour for 72 hours using a plate reader. Maximum cell density was calculated as:

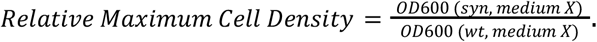

### Statistical Analysis

All experiments were performed in three biological replicates and two technical replicates in at least three independent days. Statistical significance was assessed using ANOVA, with a significance threshold of p < 0.05.

## Code Availability

The promoter design software developed in this study is released under the GPL-3.0 license and can be accessed via GitHub at https://github.com/GeorgievLab/design_of_promoter_logic. A browser-based implementation is also available at https://www.kky.zcu.cz/en/Tools/placeback.

## Acknowledgment

This work was carried out with the support of ELIXIR CZ Research Infrastructure (ID LM2023055, MEYS CR). Computational resources were provided by the e-INFRA CZ project (ID:90254), supported by the Ministry of Education, Youth and Sports of the Czech Republic.

We gratefully acknowledge the resources and know-how provided by XENO Cells Innovations s.r.o., as well as valuable consultations with its employees. We thank Filip Jani for developing the server framework and preparing the initial version of the front-end interface, Jiri Fatka for consultations on online application deployment, and Lucie Houdová for her support and guidance.

## Supplementary Information

### Data

Recommended promoters for recombination with tetO: https://github.com/GeorgievLab/design_of_promoter_logic/blob/main/data/promoters/promoters_for_tetO_recombination.json

**Supplementary Figure 1:**
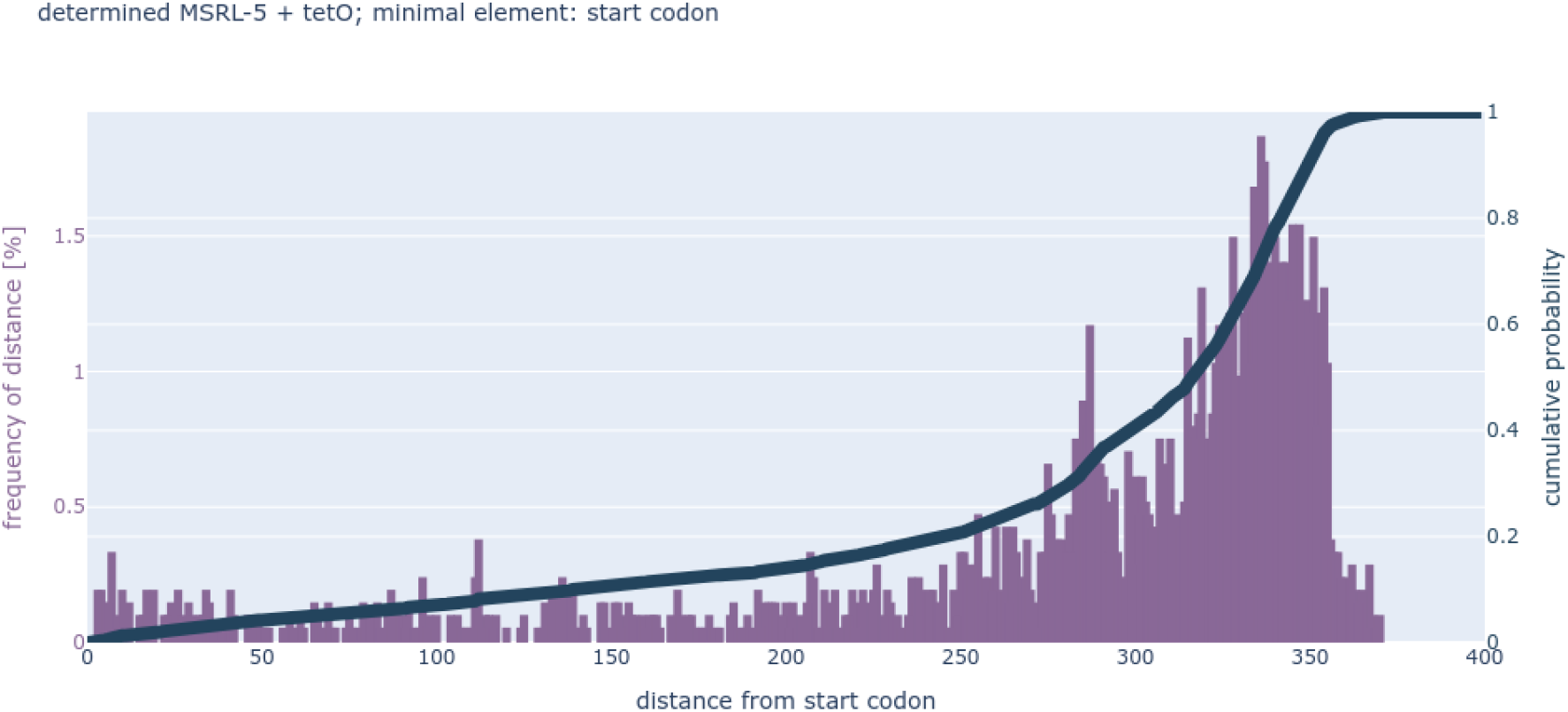
Distribution of tetO placement distances relative to the start codon in recommended promoters when models with an MSRL of 5bp were selected.

**Supplementary Figure 2:**
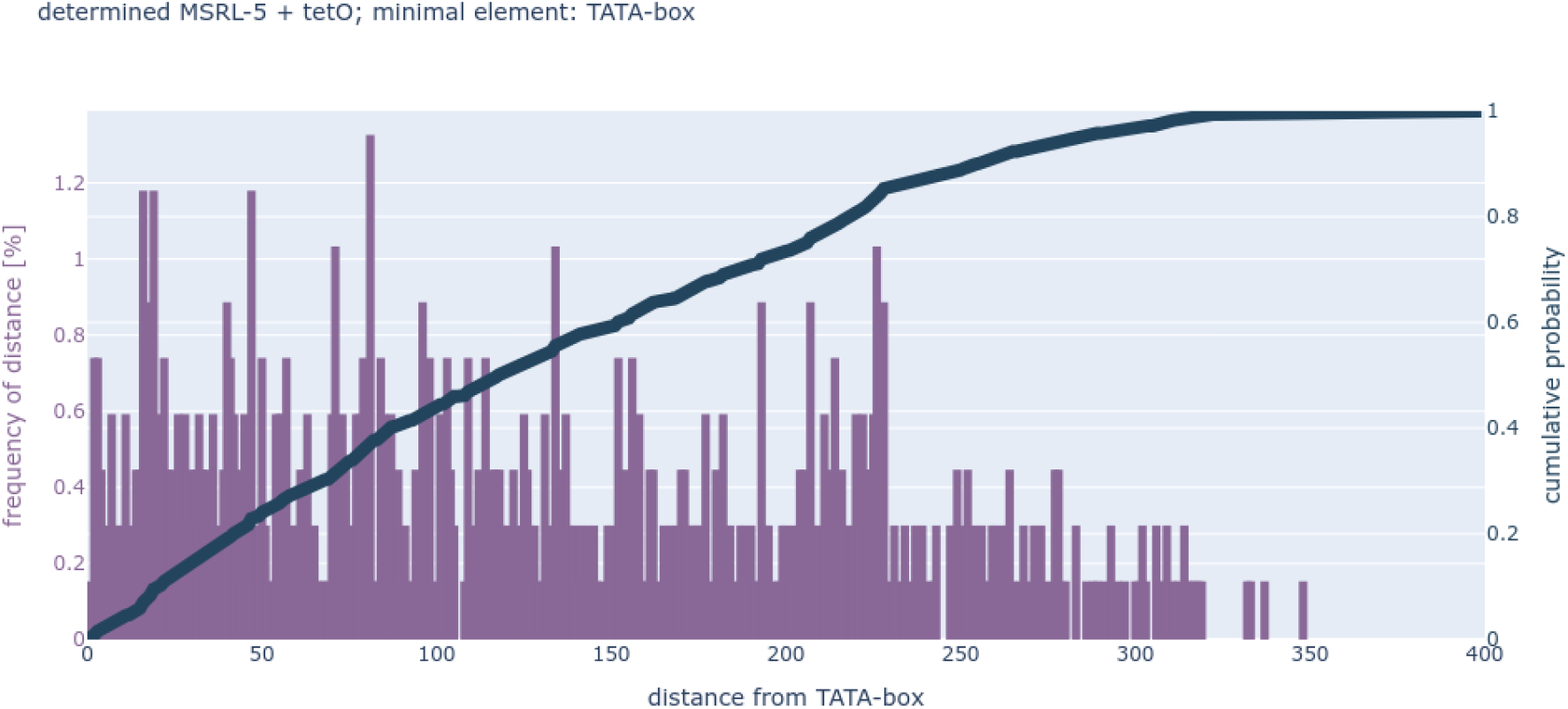
Distribution of tetO placement distances relative to the TATA-box in recommended promoters when models with an MSRL of 5bp were selected.

**Supplementary Figure 3:**
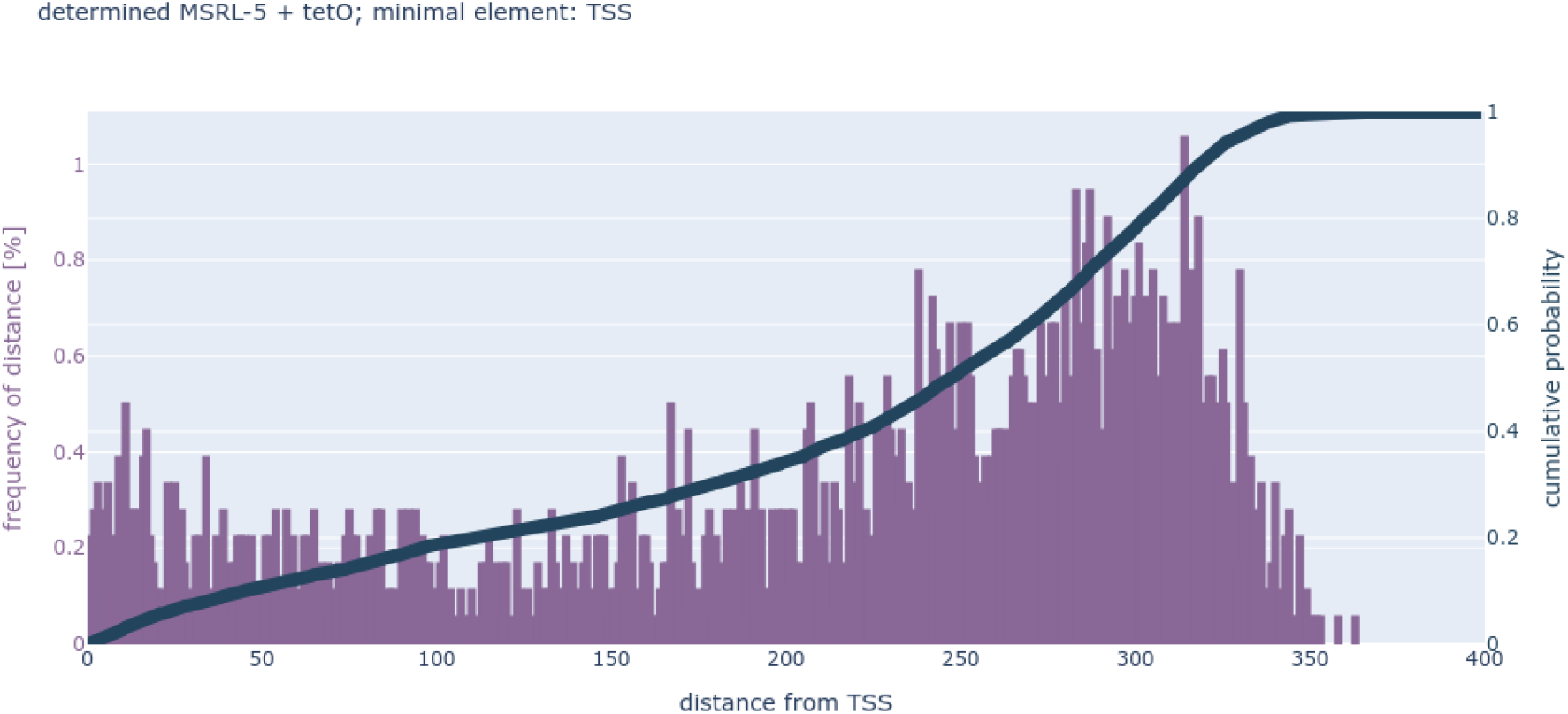
Distribution of tetO placement distances relative to the TSS in recommended promoters when models with an MSRL of 5bp were selected.

**Supplementary Figure 4:**
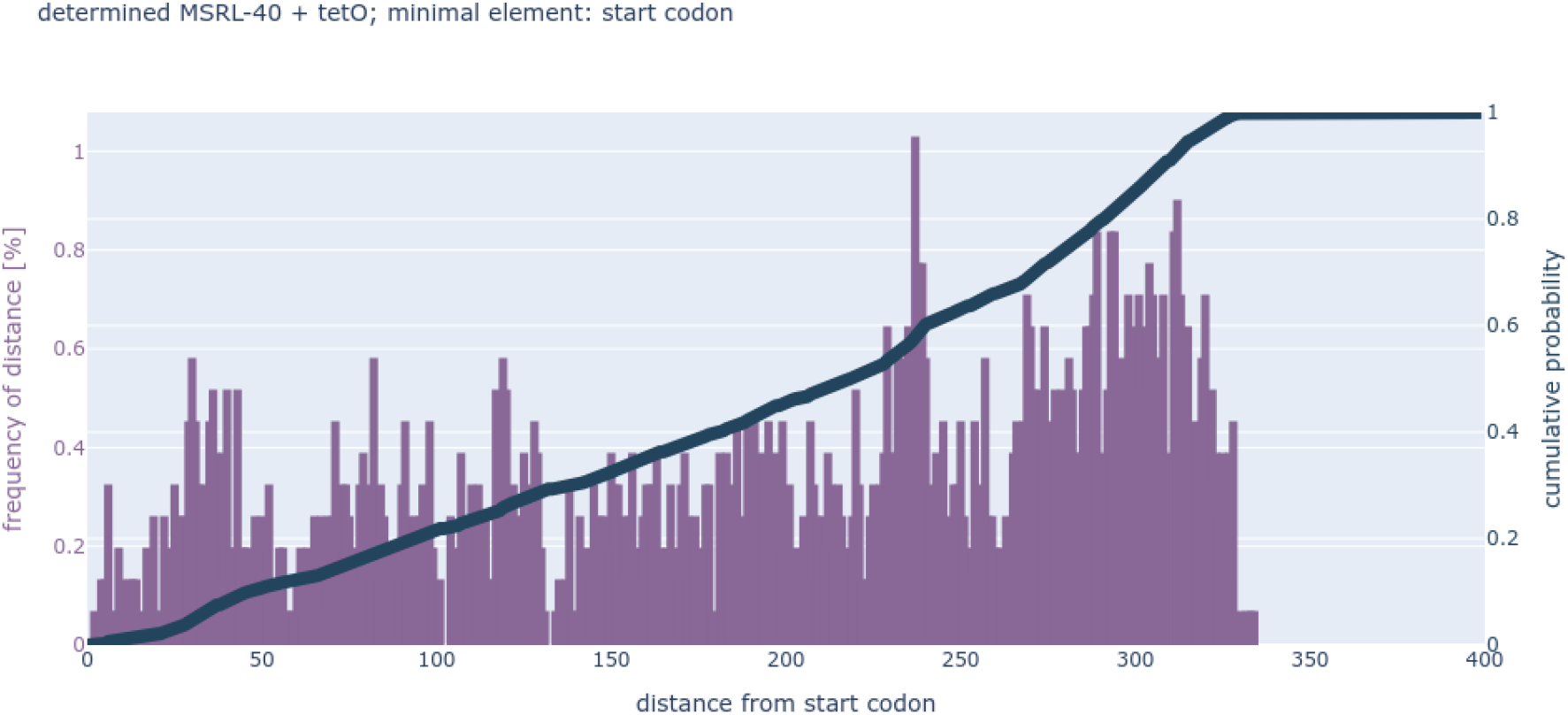
Distribution of tetO placement distances relative to the start codon in recommended promoters when models with an MSRL of 40bp were selected.

**Supplementary Figure 5:**
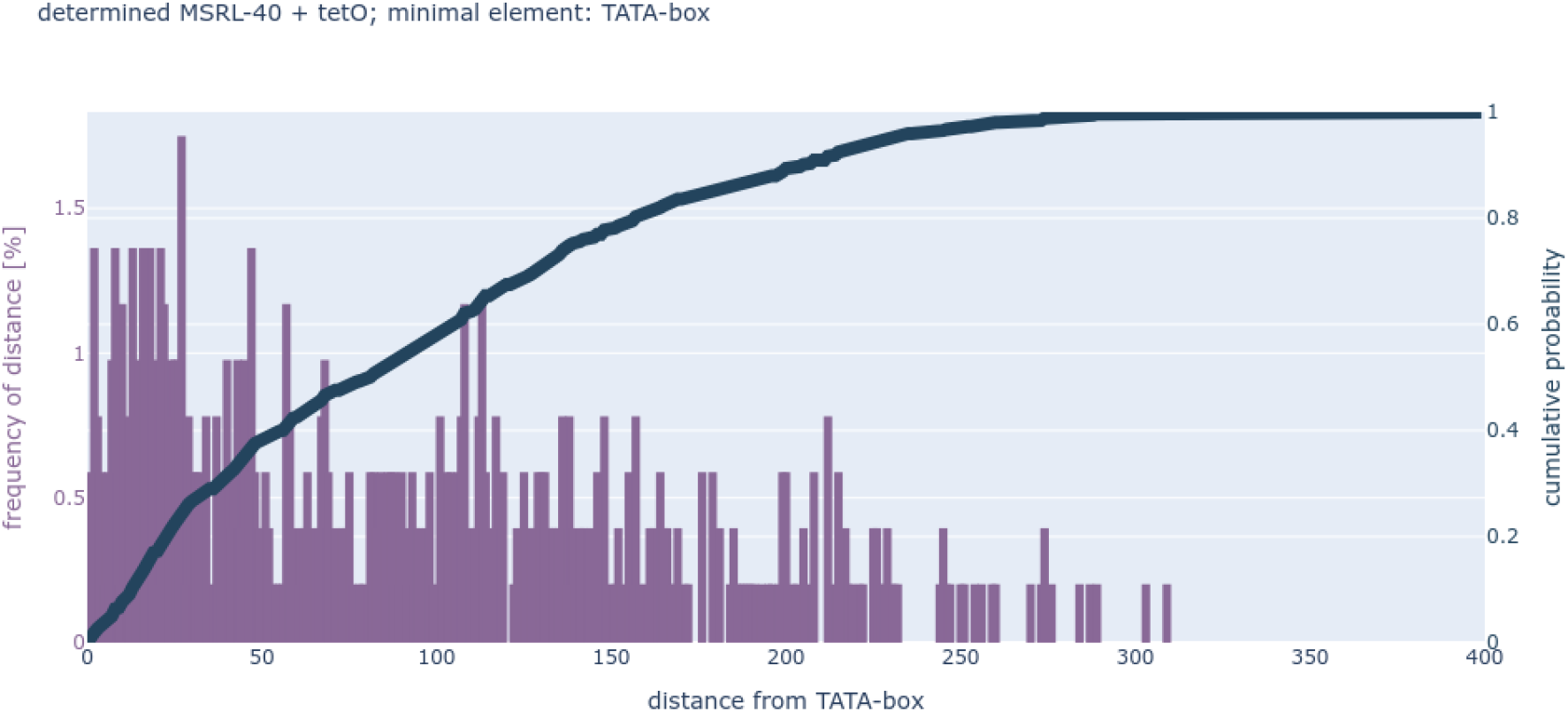
Distribution of tetO placement distances relative to the TATA-box in recommended promoters when models with an MSRL of 40bp were selected.

**Supplementary Figure 6:**
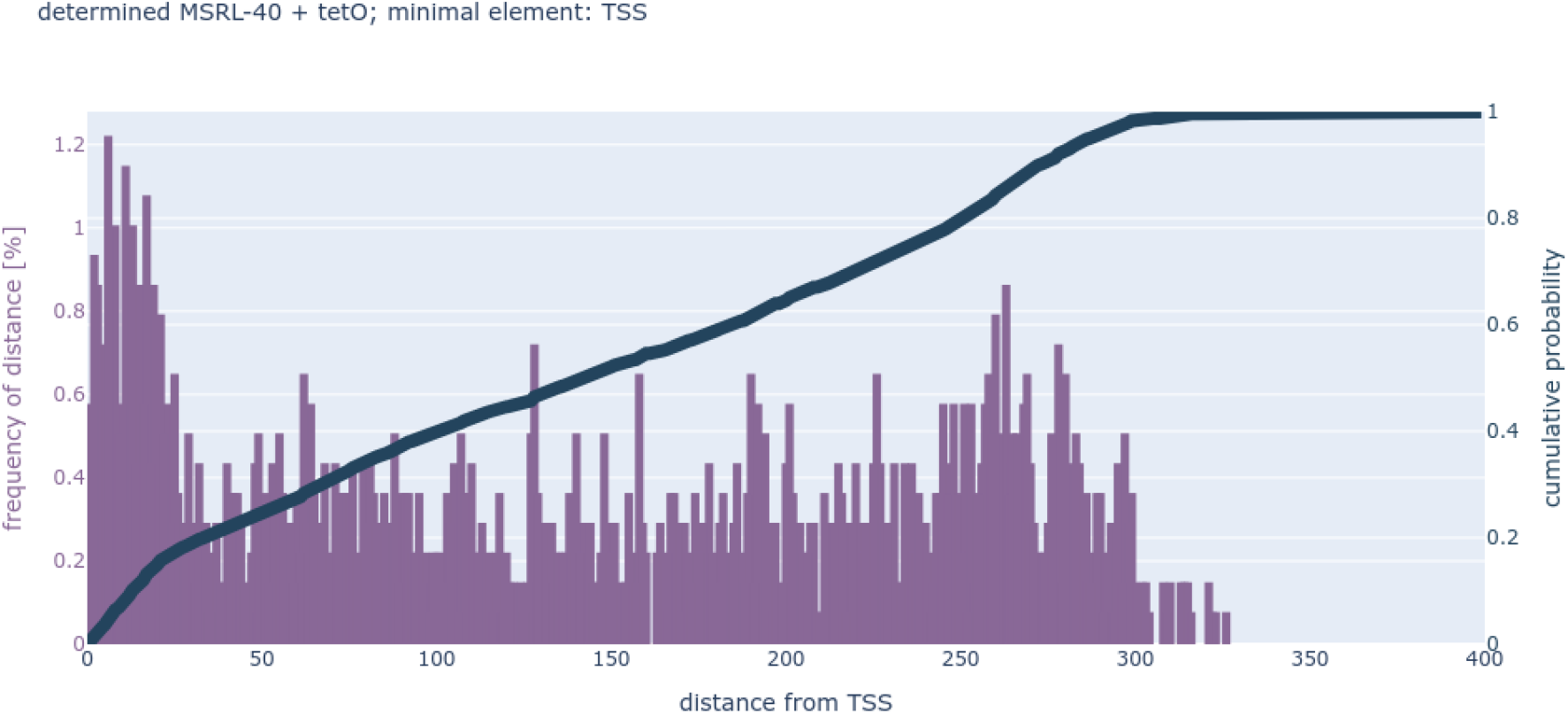
Distribution of tetO placement distances relative to the TSS in recommended promoters when models with an MSRL of 40bp were selected.

**Supplementary Figure 7:**
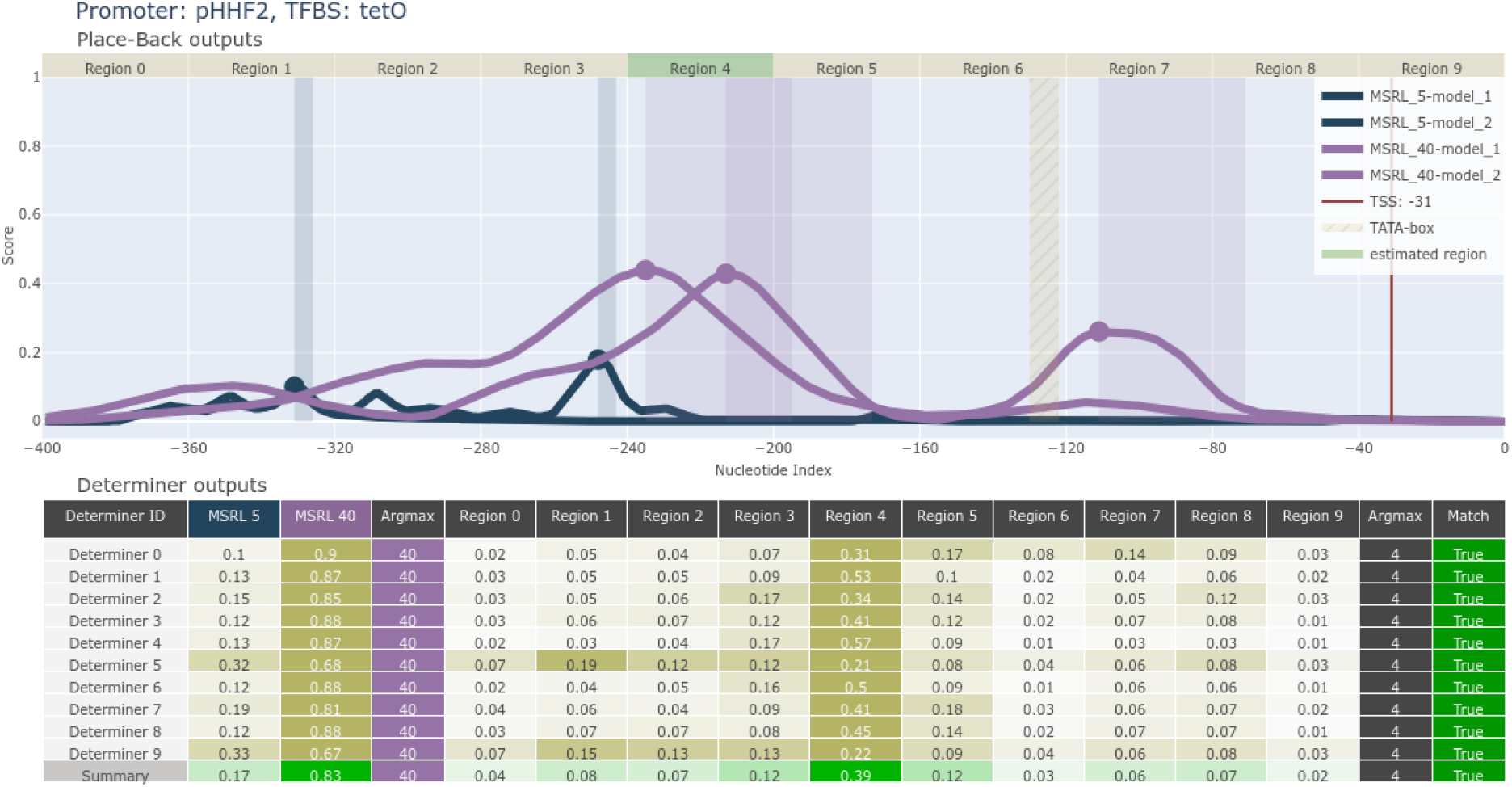
Acomplete output from the two-stage ANN system for recombination of the wild-type pHHF2 promoter with tetO. Selected tetO placement for laboratory experiment was from −235 to −195 in the promoter.

**Supplementary Figure 8:**
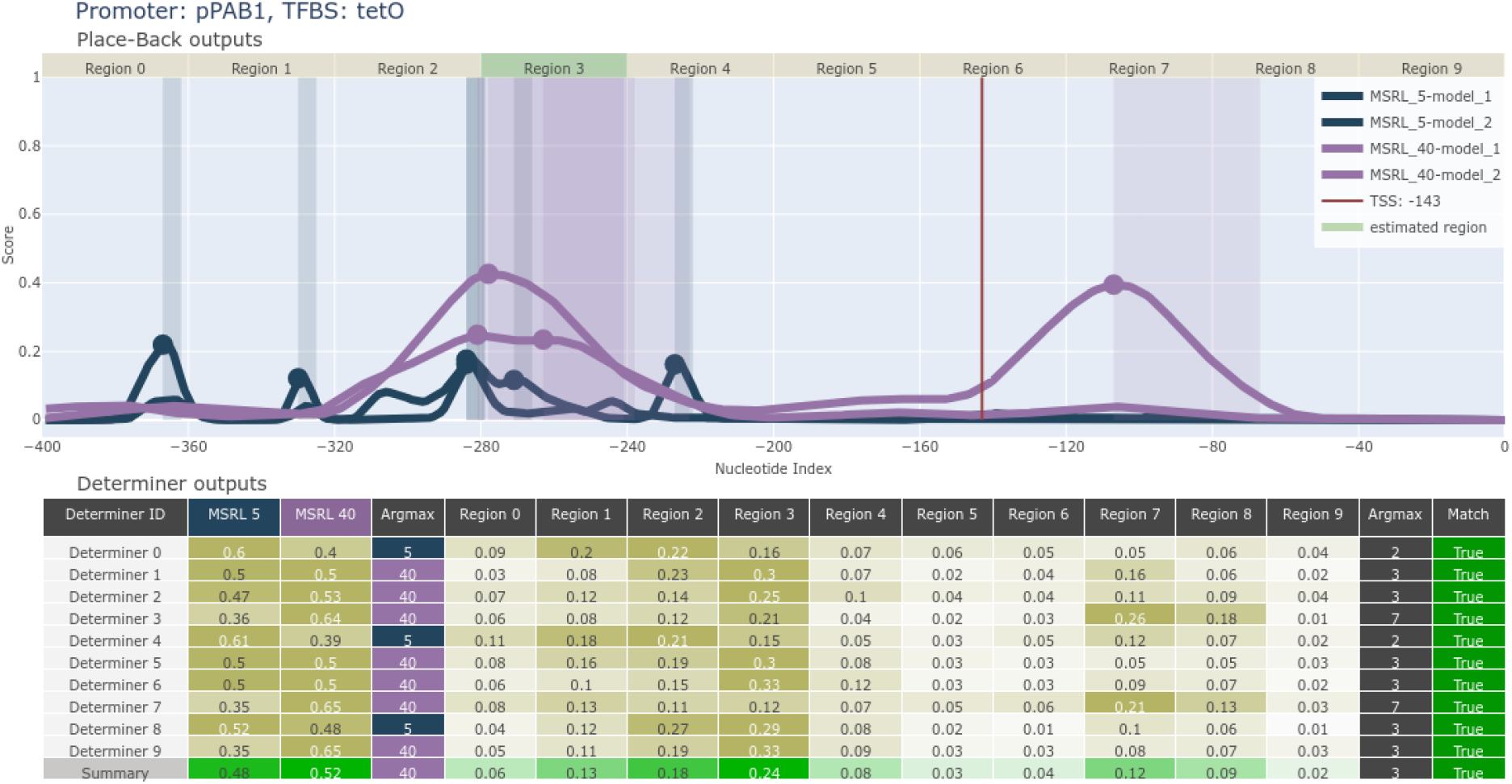
Acomplete output from the two-stage ANN system for recombination of the wild-type pPAB1 promoter with tetO. Selected tetO placement for laboratory experiment was from −278 to −238 in the promoter.

**Supplementary Figure 9:**
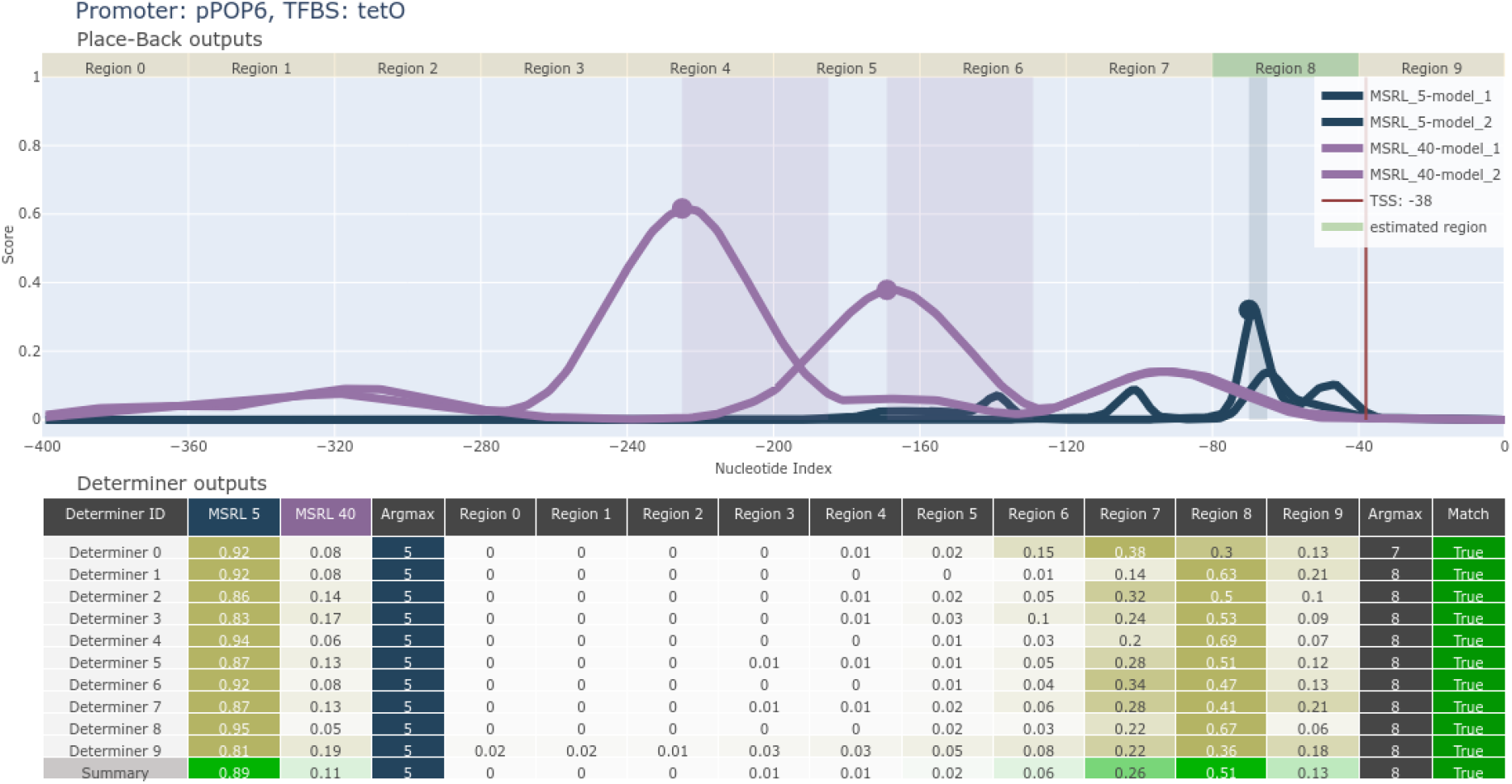
A complete output from the two-stage ANN system for recombination of the wild-type pPOP6 promoter with tetO. Selected tetO placement for laboratory experiment was from −70 to −65 in the promoter.

**Supplementary Figure 10:**
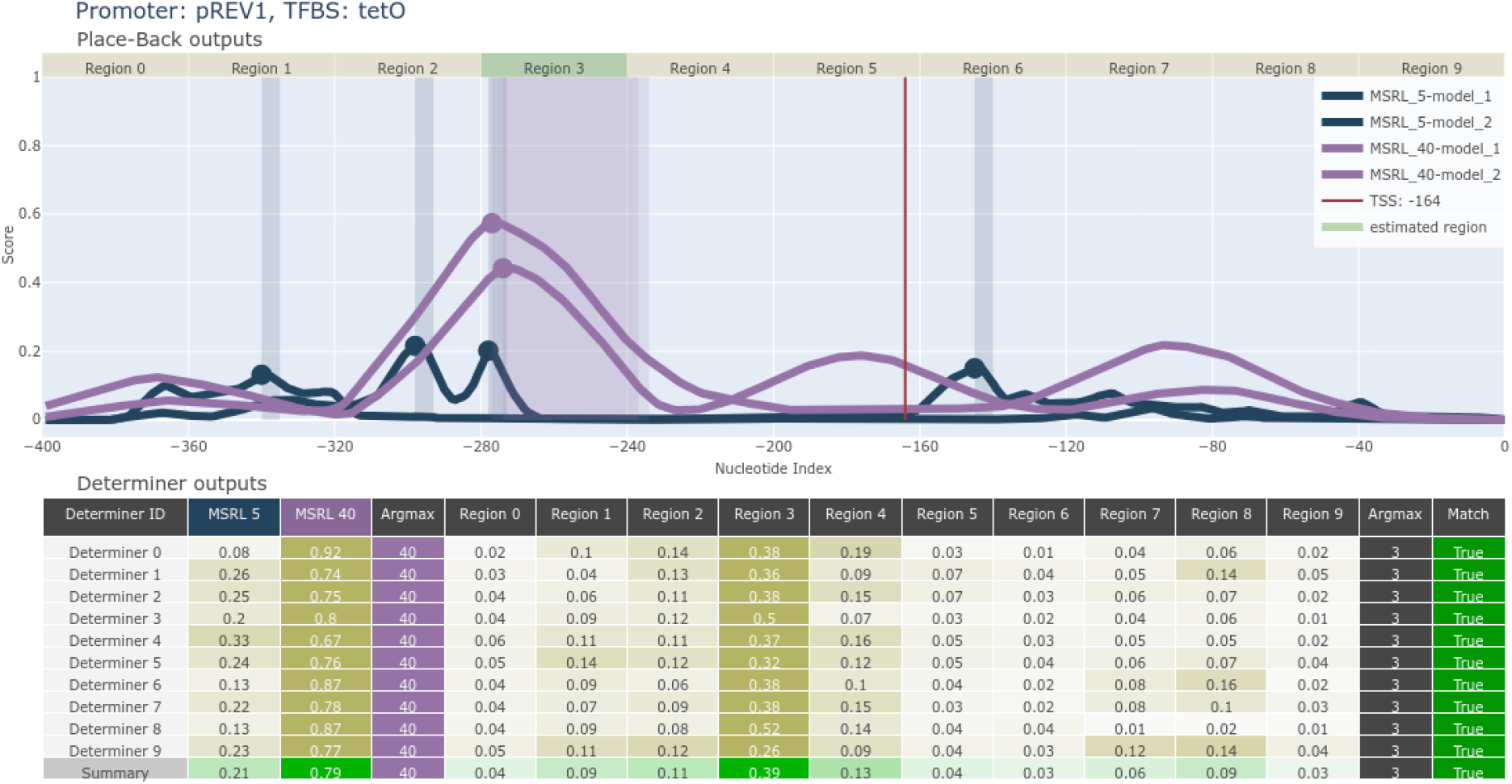
A complete output from the two-stage ANN system for recombination of the wild-type pREV1 promoter with tetO. Selected tetO placement for laboratory experiment was from −277 to −237 in the promoter.

**Supplementary Figure 11:**
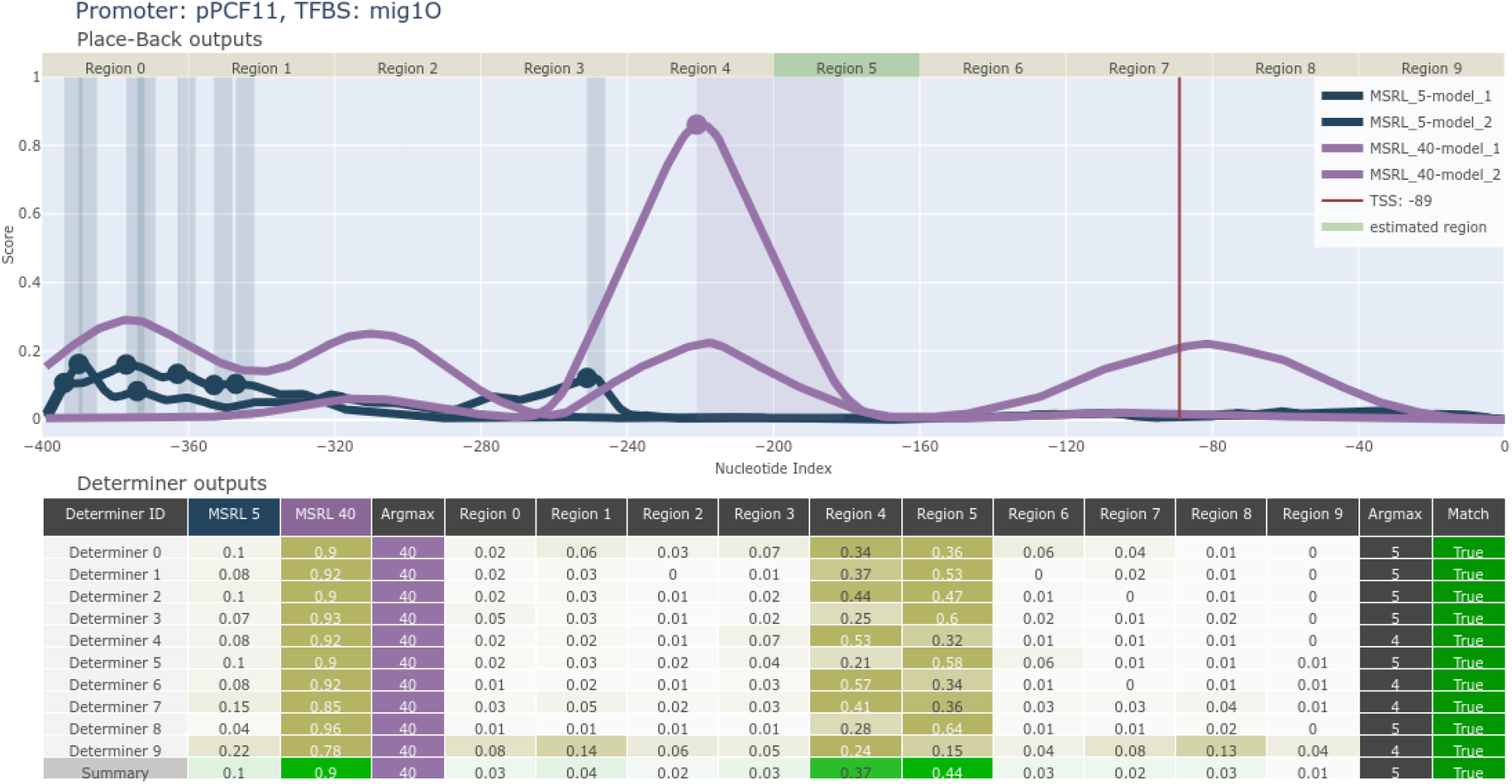
A complete output from the two-stage ANN system for recombination of the wild-type pREV1 promoter with tetO. Selected tetO placement for laboratory experiment was from −277 to −237 in the promoter.

## Notes

### Competing Interest Statement

The authors have declared no competing interest.

